# Wide-field fluorescence lifetime imaging of single molecules with a gated single-photon camera

**DOI:** 10.1101/2024.09.17.613468

**Authors:** Nathan Ronceray, Salim Bennani, Marianna Fanouria Mitsioni, Nicole Siegel, Maria J. Marcaida, Claudio Bruschini, Edoardo Charbon, Rahul Roy, Matteo Dal Peraro, Guillermo P. Acuna, Aleksandra Radenovic

## Abstract

Fluorescence lifetime imaging microscopy (FLIM) is a powerful tool to discriminate fluorescent molecules or probe their nanoscale environment. Traditionally, FLIM uses time-correlated single-photon counting (TCSPC), which is precise but intrinsically low-throughput due to its dependence on point detectors. Although time-gated cameras have demonstrated the potential for high-throughput FLIM in bright samples with dense labeling, their use in single-molecule microscopy has not been explored extensively. Here, we report fast and accurate single-molecule FLIM with a commercial time-gated single-photon camera. Our optimized acquisition scheme achieves single-molecule lifetime measurements with a precision only about three times less than TCSPC, while imaging with a large number of pixels (512×512) allowing for the spatial multiplexing of over 3000 molecules. With this approach, we demonstrate parallelized lifetime measurements of large numbers of labeled pore-forming proteins on supported lipid bilayers, and temporal single-molecule Förster resonance energy transfer measurements at 5-25 Hz. This method holds considerable promise for the advancement of multi-target single-molecule localization microscopy and biopolymer sequencing.

## Introduction

Fluorescence lifetime measurements report the nanosecond-scale delay between an excitation laser pulse and the fluorescence emitted by a molecule. The decay of fluorescence after excitation typically follows an exponential distribution, characterized by a time constant known as the fluorescence excited state lifetime, *τ*. Similarly to the emission spectrum, the lifetime contains unique information about the electronic structure of the emitter, with common dye lifetimes ranging from 0.3 to 5 ns^1,2^. Furthermore, fluorescence lifetimes are sensitive to the surrounding environment, particularly through solvent effects and energy transfer processes^1^. Lifetime measurements enable the monitoring of biochemical reactions^3^ and can be used to determine (sub)nanometer-scale distances via Förster resonance energy transfer (FRET), allowing the detection of molecular interactions or conformational changes in proteins^4^. Fluorescence lifetimes can also encode information about the axial localization of emitters relative to conductive substrates^5,6^. Typically, FLIM relies on TCSPC integrated on a confocal microscope with a point detector that records photon arrival times. Super-resolved variants of scanning confocal microscopy have been demonstrated with lifetime-resolved MINFLUX^7,8^ and RASTMIN^9^, but none of these techniques addresses several targets in parallel, making them low-throughput despite their unmatched localization precision.

Conversely, wide-field FLIM enables assigning lifetime values to multiple targets using parallel detection with a detector array, offering much higher throughput. Wide-field TCSPC has been investigated^10^ even at the single-molecule level, but sensors that enable it suffer from constraints on sample brightness and a very low photon detection efficiency (PDE)^11,12^. Rather than recording photon arrival times, wide-field FLIM can be achieved by acquiring time-delayed images on a camera, which allows acquiring only specific temporal windows of the fluorescence decay from the sample. Although this delay can be obtained spatially^13,14^, it is typically achieved by using a time gate that collects photons exclusively within a specific time window after excitation. In most implementations, this time gate can be moved relative to the excitation pulse, allowing accurate sampling of the fluorescence decay. Gate positioning with picosecond-scale resolution can be achieved using intensified charge-coupled device cameras^15,16^ or by using a Pockels cell in the emission path^17,18^. The latter approach proved very efficient at fast recording of neural activity^19^, but showed imprecise results in the context of single-molecule imaging^17,18^.

Single-photon avalanche diode (SPAD) arrays have emerged as readout-noise-free detectors with a good PDE^20^. Although most of the fluorescence lifetime studies based on SPAD arrays used small asynchronous arrays (ranging from 20 to 400 detectors) with TCSPC capabilities^21–24^, wide-field FLIM has also been demonstrated using synchronous time-gated SPAD cameras with nearly a million pixels^25^. In such SPAD cameras, the gating capability is monolithically integrated in the sensor, unlike intensifier- or Pockels cell-based systems, thus offering better robustness and excellent temporal resolution.

Considerable efforts have been dedicated to establishing a palette of techniques for separating multi-exponential decays from a diffraction-limited spot^1,2^ including the case of gated SPAD cameras^26^. Such techniques are generally photon-greedy and thus poorly suited for single-molecule lifetime measurements. However, single-molecule fluorescence decays tend to be simpler as a result of their well-defined electronic properties and local environment. Indeed, single-molecule decays are generally mono-exponential, unlike those of molecular ensembles where two or more exponential components can be found due to the presence of multiple dye populations in different local environments within the same imaged volume^11,27^. Although they have simpler decay shapes, single molecules present constraints because of their limited brightness and low photon budget. Hence, it is not surprising that single-molecule lifetime imaging has not been thoroughly investigated with gated SPAD cameras. In this work, we use the SPAD512^2^ camera^25,28^ that has a PDE of 30-40% when using microlens arrays, which allows for single-molecule imaging^29^. With this sensor, we revisit a gated imaging scheme introduced 35 years ago^30^ and demonstrate its potential for single-molecule fluorescence lifetime imaging microscopy (smFLIM).

## Results

### Single molecule (sm)FLIM scheme and setup

We designed a fluorescence microscopy setup using the SPAD camera to image molecules with total internal reflection illumination (Fig. 1a, details in Methods). The time gate is synchronized with the ~ 50 ps pulsed supercontinuum light source used for wide-field illumination at a repetition rate of 26 MHz. The gate profile can be approximated by a square function with a rise time of ~ 200 ps and a controllable gate width *W* in the range of 6 − 12 ns (Fig. S10a). Controlling the relative delay of the gate opening with respect to the excitation pulse, we implement the so-called rapid lifetime determination scheme^30^ consisting of alternating two gate positions. The first gate, positioned at the very beginning of the decay, collects nearly all photons from the sample, and the second gate is offset by a delay *T* to reject the early emitted photons. We show in Figure 1b typical images of molecules with the full fluorescence signal in the first gate and a dimmer version of the image in the second gate. Mathematically, this 2-gate scheme enables retrieving the fluorescence lifetimes of molecules exhibiting monoexponential decay. Indeed, a molecule with unknown lifetime *τ* emitting a signal of *S* photons in the first gate, imaged with a background *B*_*o*_ will yield *N*_*o*_ = *S* + *B*_*o*_ photons in the first gate and *N*_1_ = *Se*^*−T/τ*^ + *B*_1_ in the second gate (Fig. 1c). The fluorescence lifetime *τ* is thus obtained as:

**Fig. 1:**
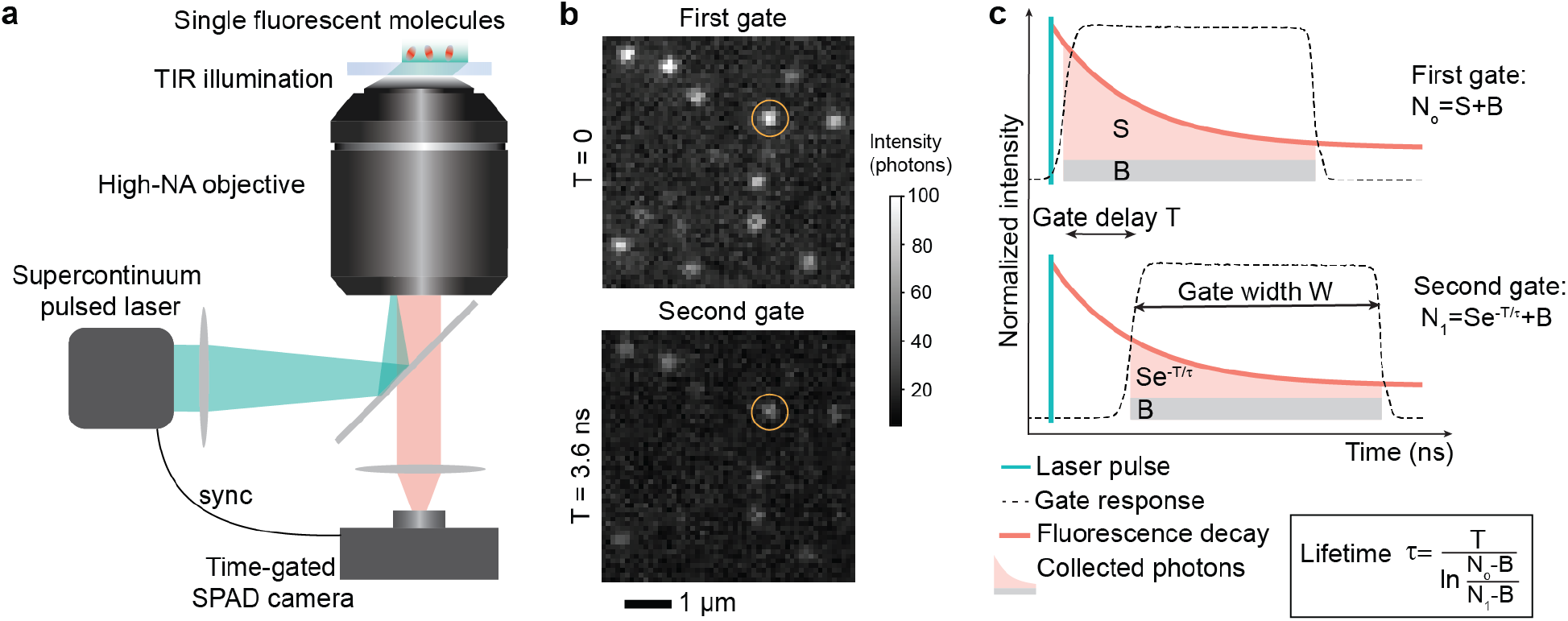
Single molecule FLIM setup and acquisition scheme. **a**, Sketch of the experimental setup where a picosecond-pulsed green laser beam undergoes total internal reflection at the sample coverslip to excite single fluorescent molecules close to its surface. The emitted fluorescence is collected to form an image on the camera chip, which is synchronized with the excitation laser. **b**, Typical images of single molecules obtained from 20 ms integration time for the first gate capturing the full fluorescence decay (top) and the second gate capturing only late photons reaching the camera after a gate delay *T*. The same molecule is shown by yellow circles in the two cases. **c**, Sketch of the time-gated imaging scheme that enables retrieving single-molecule fluorescence lifetimes. In the first gate (top) the *N*_*o*_ detected photons can be decomposed into single-molecule fluorescent signal (*S*) and background (*B*). In the second gate (bottom) early fluorescence is rejected and fewer photons are measured (*N*_1_).

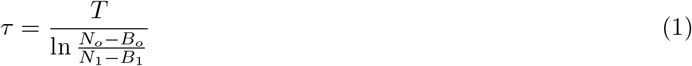

The background *B*_*o*_ and *B*_1_ measured in the first and second gate may be different in the eventuality of a fluorescent component of the background. In the following, we apply this scheme to highly multiplexed single-molecule lifetime measurements. Although the 2-gate scheme employed here cannot outperform standard TCSPC-based FLIM approaches in terms of photon efficiency, it enables massive multiplexing by measuring a large number of lifetimes at once from a single field of view.

We assumed *B*_*o*_ = *B*_1_ = *B* for simplicity on this sketch but in the general case we use *B*_*o*_ ≠ *B*_1_ = *B*. The gate response is mapped in more details in Figure S10.

### High-throughput smFLIM on bacterial pore-forming proteins on lipid membranes

Our setup enables the imaging of thousands of individual molecules within a 51 × 51 μm^2^ field of view, with a 100 nm pixel size at the sample. To demonstrate the high-throughput capability of our system, we imaged labeled aerolysin proteins embedded in supported lipid bilayers (Fig. 2b). Aerolysin is a *β*-barrel pore-forming toxin that assembles into oligomeric structures, forming transmembrane nanopores widely used in biosensing^31^. In our experiments, mixed labeled and unlabeled aerolysin monomers are initially present in the incubation solution and progressively integrate into the membrane, forming stable singly labeled heptameric assemblies within the lipid bilayer. This approach allows for control over the molecular density by adjusting the concentration of the solution and the incubation time. Most of our images were acquired at a density of roughly one labeled aerolysin pore complex per μm^2^ (Fig. 2a). Detailed sample preparation methods are provided in the Methods section. The intensity traces of the first gate *N*_*o*_(*t*) and the second gate *N*_1_(*t*) were extracted for each of the ~ 3000 fixed molecules in the field of view, allowing the application of stringent selection criteria to keep the pure single-molecule traces. For this, we used a python-based analysis pipeline relying on molecule detection (yellow squares in Fig. 2a) followed by filtering based on the identification of single-step photobleaching (red squares, see Methods) to ensure that the detected spot corresponds to a single molecule. Molecular lifetimes were then extracted by applying equation 1 to the time-averaged first and second gate signals. This enabled obtaining well-sampled lifetime histograms in under a minute, considerably speeding up the acquisition of lifetime statistics compared with standard confocal approaches. We show in Figure 2c histograms for proteins labeled with various dyes (LD555, Cy3B, and AF488) exhibiting different lifetimes and wavelengths. The measured lifetime values are presented in Table S1.

**Fig. 2:**
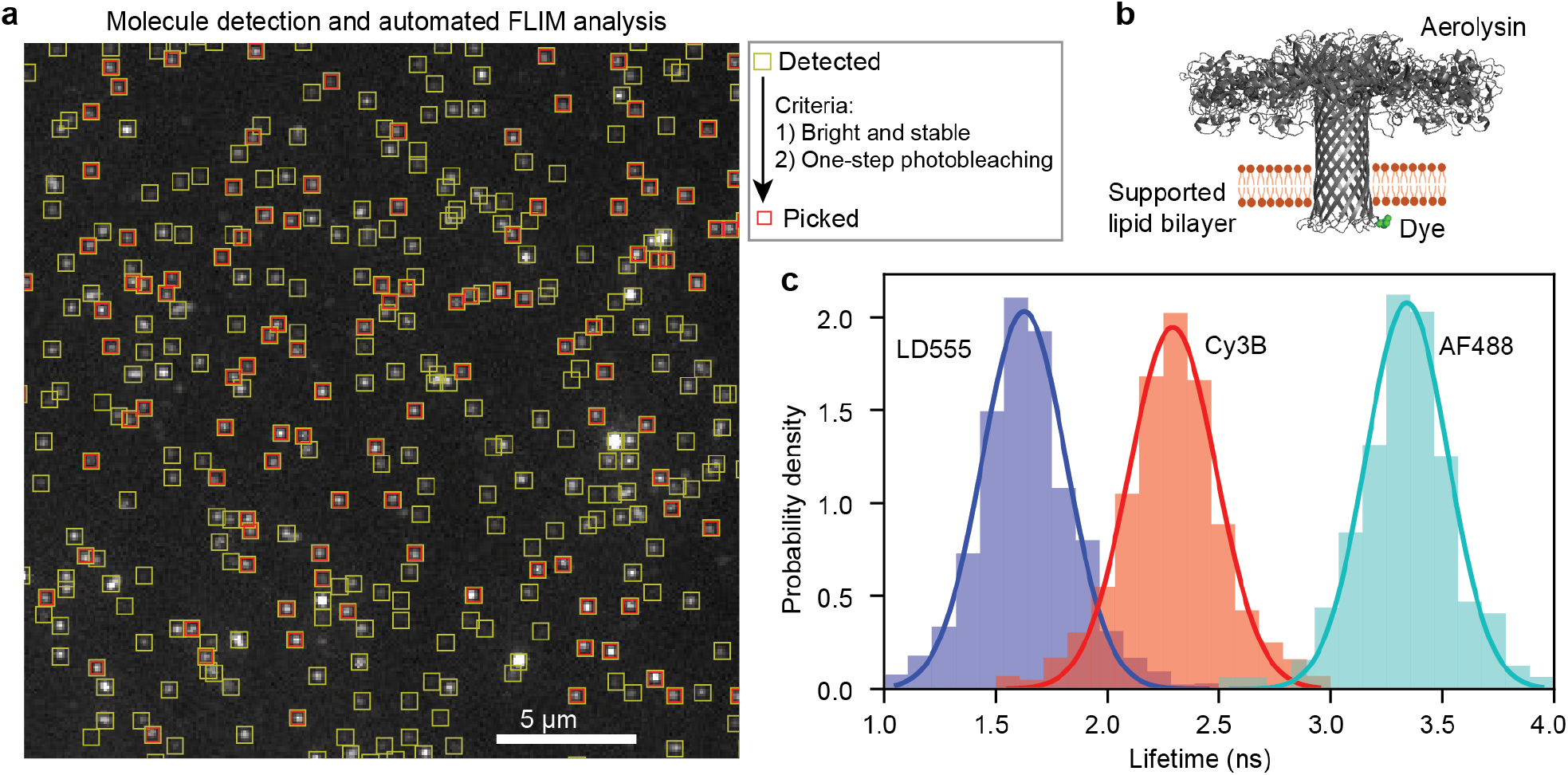
Spatially multiplexed single-molecule FLIM of labeled membrane proteins. **a**, Image of Cy3B-labeled aerolysin proteins in a supported lipid bilayer, showing detected molecules (yellow square) and picked molecules (red square) based on their suitability for lifetime assignment. **b**, Structure of the pore-forming aerolysin protein and position of the label. **c** Lifetime distributions obtained from 5 fields of view for LD555 (*n* = 1643), Cy3B (*n* = 2380) and AF488 (*n* = 301) dyes, respectively. For each distribution the total acquisition time is less than one minute. The gate delay is 2.5 ns for LD555 and Cy3B and 3 ns for AF488.

We verified that the gate delay value did not cause any systematic bias in the estimation of the lifetime distribution (Fig. S1a). We then estimated the throughput improvement of our scheme over sequential scanning with a point detector. Considering that all molecules are excited together, the acquisition must be long enough to capture all traces. We find that with a median photobleaching time of approximately one second, acquiring images for 10 seconds ensures that all photobleaching events are captured (Fig. S1b). Thus, while point scanning would require 3000 seconds to sequentially image the photobleaching of 3000 molecules, our wide-field scheme takes only 10 seconds. This 300-fold acceleration of single-molecule FLIM measurements opens the door to screening large populations and identifying potentially rare events among fluorescent traces. While these results demonstrate the potential of our scheme for *static* lifetime measurements, they rely on large photon budgets on the order of 10^4^ (see Fig. S1c) which are not limited by shot noise.

### Optimizing smFLIM

We now turn to the photon efficiency of our approach with the goal of quantifying its potential for *dynamic* lifetime measurements. Due to the low brightness levels characteristic of single-molecule imaging, optimizing the acquisition scheme is essential. For the ability to distinguish different lifetimes in a given sample, it is important to minimize the spread of the measured lifetime values of a given molecule because it will establish the achievable temporal resolution of lifetime measurements. This spread originates mainly from the shot-noise-induced error in the lifetime estimate. The standard metric to quantify the spread of lifetimes obtained through FLIM techniques is the *F*-value^20,32^ that is defined as:

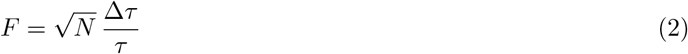

Where *N* is the number of available photons used to report a lifetime estimate (*N* = 2 × *N*_*o*_ in our scheme), *τ* and Δ*τ* are the mean and standard deviation of the lifetime estimate distribution, respectively. For example, with *N* = 1000 photons available, the relative error on the lifetime is 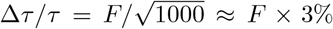. TCSPC theoretically achieves *F* = 1 in background-free conditions with a large number of channels^33^, which means perfect photon efficiency of lifetime measurements, but SPAD array-based TCSPC was limited to *F* ≥ 1.5 so far^20^. Gated imaging cannot reach perfect photon efficiency as it relies on rejecting photons. Our 2-gate scheme minimizes photon rejection compared with multi-gate schemes but still rejects a fraction(*N*_*o*_ − *N*_1_)*/N* = (1 − *e*^*−T/τ*^) × *S/N* of the available photons.

To evaluate the *F*-value of our scheme, we imaged fluorescent beads and systematically varied the main parameters of our system: the normalized gate delay *u* = *T/τ*_ref_ and the signal-to-background ratio (SBR) *S/B* (Fig. 3a). To modulate the background level, we used the reflection of a thermal white-light illumination of various intensities superimposed with the pulsed laser illumination of fixed intensity. The beads’ ground-truth fluorescence lifetime *τ*_ref_ was measured through a multi-gate approach (details in Figure S2). We then examine the variations of the lifetime statistics as obtained from equation 1 depending on the normalized gate delay *u* and the SBR, and evaluate *F*-values through equation 2 (Fig. 3b). The optimal gate delay depends strongly on the SBR: for a large SBR (blue curve), the optimal value of *T* is slightly above 2*τ*, in accordance with previous studies relying on Monte Carlo simulations^34,35^. The case of low SBR (red curve) differs in the optimum: the optimal value of *T* is slightly above *τ*. The results are in full agreement with the analytical calculation obtained by Heeg^36^, which we adapted and extended in the Supplementary Information and in Figure S7. In Figure 3c, the optimal gate delay (top) and resulting *F*-value (bottom) are presented. The region highlighted in gray corresponds to the regime relevant to single-molecule imaging, in which the strong variation of the optimal gate delay shows the importance of carefully optimizing imaging parameters for such conditions. The resulting *F* values in this region are in the range of 2 − 6, indicating moderate loss of photon efficiency compared to TCSPC. To obtain a given relative lifetime uncertainty Δ*τ/τ* the integration time should be set such that 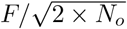 is small enough.

**Fig. 3:**
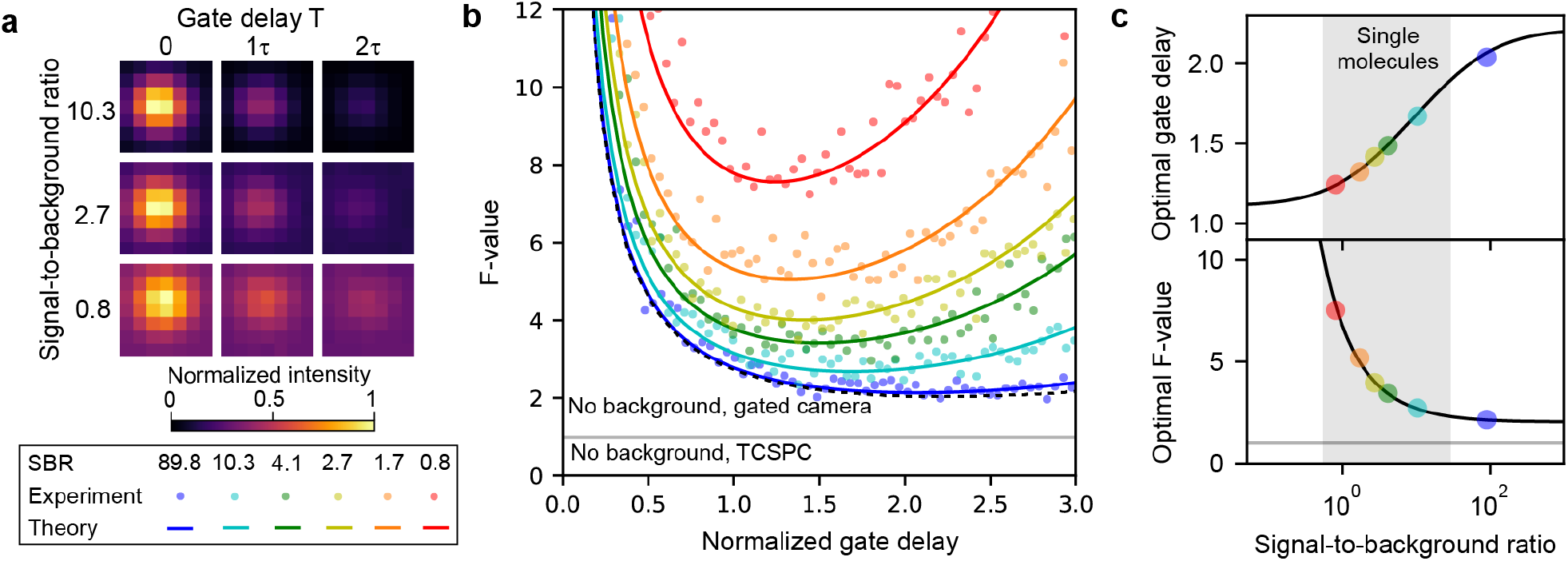
Optimization of the acquisition scheme. **a**, Experimentally varying parameters using fluorescent bead imaging with controlled gate delay *T* and controlled background resulting in different SBR values. **b**, Experimental *F*-value are compared with the theoretically expected formula 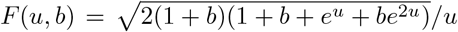 where *b* = 1*/*SBR and *u* = *T/τ* is the normalized gate delay. **c**, Optimal gate delay (top) and minimum *F*-value at this optimal delay value (bottom) as a function of the emitter ‘s SBR. Solid lines are numerically evaluated from deriving the minimum of the *F*-value formula, and markers indicate experiments.

### Single-molecule FRET and dynamic smFLIM measurements

Having shown the spatial multiplexing power of our scheme and identified the optimal operation scheme, we now demonstrate its applicability to monitoring lifetime changes in real time, as exemplified by changes in single-molecule FRET efficiency^37^. We imaged DNA origami labeled with a donor dye (Cy3B) and an acceptor dye (Alexa 647) within the FRET range (*d* ≈ 5.7 nm, details in Methods). We used a 600 nm shortpass filter in the emission path to image donor molecules only, and quantified FRET through their lifetimes. The sample contained origamis with only a donor and some with a donor-acceptor pair, resulting in distinct lifetime values *τ*_D_ and *τ*_DA_ (Fig. 4a-b). Extending the optimization presented in Figure 3 to the case of resolving two lifetimes, we found that the optimal gate delay value is in general close to the longest of the lifetimes in conditions used here (Fig. S8). We identified many occurrences (*n* = 7029) of traces exhibiting single-step donor photobleaching, corresponding to the criterion used in Figure 2a. As expected, single-step photobleaching lifetimes were found to contain two populations with stable lifetimes around 2.1 ns and 3.4 ns, respectively.

**Fig. 4:**
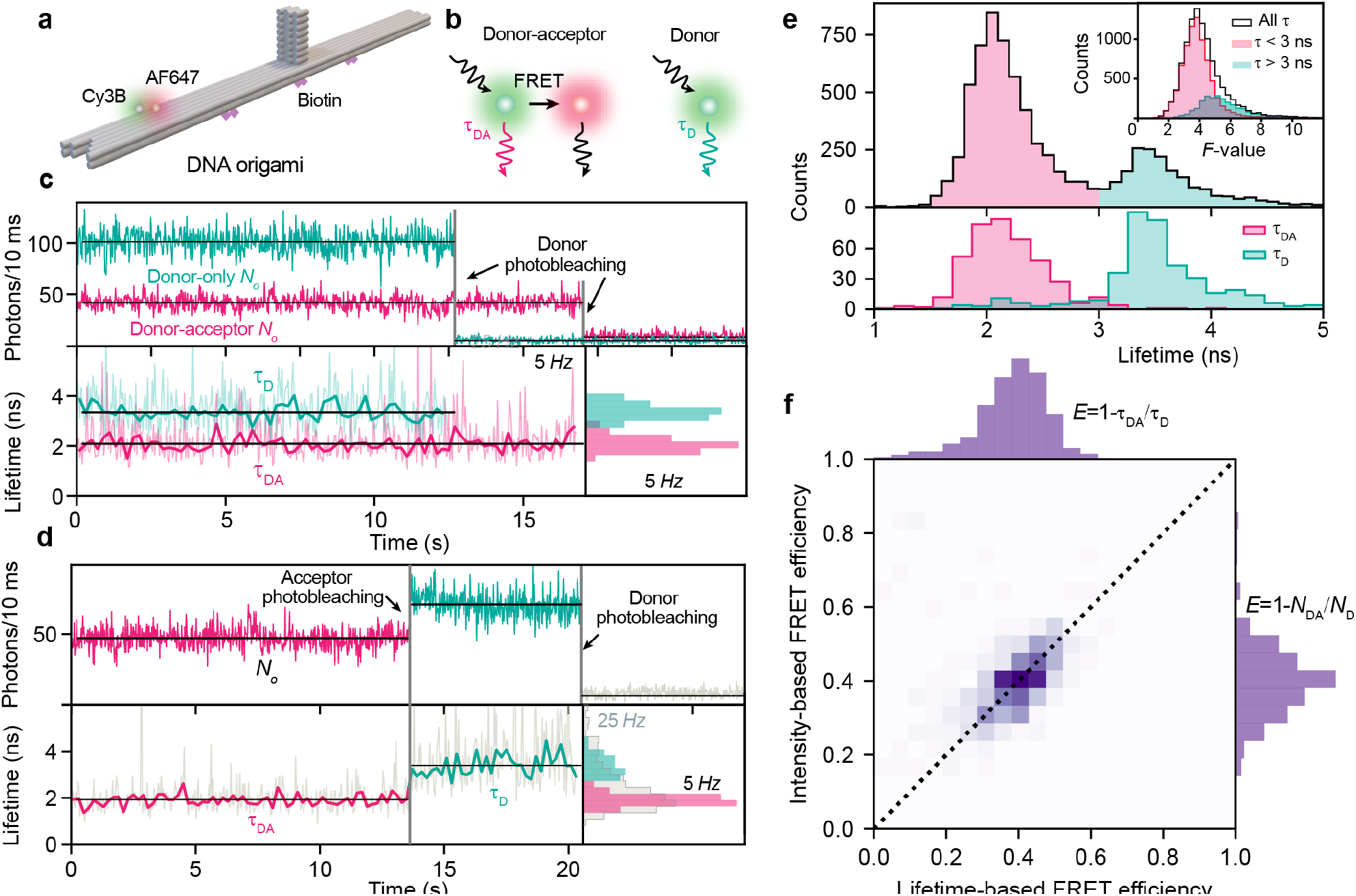
Single-molecule FRET and dynamic smFLIM. **a**, Sketch of the DNA origami design used in this experiment. **b**, Sketch of the emission process of a donor-acceptor pair in the FRET range (5.7 nm distance) where the donor has a lifetime *τ*_DA_ (left) and donor alone with lifetime *τ*_D_ (right). **c**, Top: representative intensity traces with single-step photobleaching events. Bottom: corresponding lifetime traces sampled at 5 Hz show a short donor lifetime corresponding to the FRET case, and a long donor lifetime in the absence of FRET. **d**, Representative intensity trace with successive acceptor and donor photobleaching, which enables a readout of the FRET and non-FRET donor lifetimes for the same molecule. Light traces in **c-d** sampled at 25 Hz are too noisy for a clear readout but 5 Hz sampling enables to reduce shot noise fluctuations. **e**, Lifetime distributions obtained from all events corresponding to **c** (top, *n* = 7029) and to **d** (bottom, *n* = 413). The top inset shows experimental *F*-value in the range 2 − 6. **f**, Single-molecule FRET histograms obtained through lifetime (horizontal) and intensity (vertical) show good agreement.

We examined time-resolved lifetime traces *τ* (*t*) obtained by applying equation 1 to the time-dependent first and second gate signal *N*_*o*_(*t*) and *N*_1_(*t*) (Fig. 4c-d). We find that single-molecule traces sampled at 5 Hz enable clear lifetime discrimination of the emitter types as shown in the inset histogram, but at 25 Hz, shot noise fluctuations prevent clear resolution of molecules through their lifetimes. We found an important number (*n* = 413) of intensity traces showing a stepwise increase as a result of acceptor photobleaching followed by a complete extinction due to donor photobleaching. As shown in the example trace in Figure 4d), a clear stepwise lifetime change can be resolved at 5 Hz sampling. We present in Figure 4d the full histograms of single-step donor photobleaching lifetimes (top) and acceptor-donor photobleaching traces (bottom), where precise lifetime estimates were obtained by applying equation 1 to segment-wise time averages of *N*_*o*_ and *N*_1_. Additional example traces are presented in Figure S4 and Figure S5. The top inset of Figure 4d shows the distribution of single-molecule *F*-values obtained by applying equation 2 to experimentally observed lifetime fluctuations shown in the traces of Figure 4b-c. We find *F*-values in the range of 2 − 6 in accordance with SBR conditions.

Acceptor-donor photobleaching traces as in Figure 4c enable retrieving the FRET and non-FRET lifetime values of the same molecule, yielding the single-molecule FRET efficiency^38,39^ through *E* = 1 − *τ*_DA_*/τ*_D_. We show in Figure 4e the analysis of acceptor-donor photobleaching traces to examine the FRET efficiency and compare to intensity-based lifetime measurements. We find good agreement between the lifetime-based readout of the FRET efficiency (horizontal axis) intensity-based value^38^ *E* = 1 − *N*_DA_*/N*_D_ obtained from photons in the first gate in the presence (*N*_DA_) and in the absence (*N*_D_) of acceptor. Lifetime-based FRET measurements are generally preferred over their intensity counterparts, as they are less prone to errors^40^. For instance, single-molecule lifetime measurements enable the rejection of intensity-induced artifacts, as shown in examples of two-step photobleaching traces exhibiting no lifetime changes (Fig. S6). As a result, the lifetime distribution is considerably sharper than the intensity distribution in Figure S3d. Nevertheless, lifetime measurements are more photon-greedy than intensity-based measurements, as shown by the clear intensity distinction between FRET and non-FRET at 25 Hz in Figure 4d. Thus, the smFLIM scheme offers a robust and accurate estimate of single-molecule lifetimes that complements well traditional intensity measurements in FRET applications.

## Discussion

We demonstrated wide-field single-molecule lifetime imaging that enables spatially multiplexed time-resolved measurements for monitoring dynamic lifetime changes. Our approach enables screening large numbers of molecules and identification of subpopulations exhibiting distinct molecular characteristics. We chose single-molecule FRET for the demonstration but our scheme is ideally suited for other lifetime-changing phenomena and particularly non-radiative energy transfer to bulk metal or 2D materials^5,6,41^. The latter has recently been demonstrated to enable real-time structural biology of enzymes at work through sub-nanometer axial localization^42^. Another important avenue for smFLIM is molecular discrimination for sequencing by synthesis applications^43,44^. These applications rely on the ability to resolve two dye molecules that transiently bind to an enzyme, which can be done using spectral information. Unlike lifetime decays, spectra have non-universal shapes and thus need high-resolution signal to be quantitatively measured. Ratiometric imaging^45^ offers an efficient simplification as employed in some high-throughput sequencing systems. For example, a dual view imaging system is used together with a dual-color excitation scheme, which enables resolving four different probes. Unlike these widespread approaches, our wide-field single-molecule FLIM scheme offers an optimal throughput for separating populations of molecules without splitting the single-molecule photon budget over different channels. Our approach also reduces the optics requirements which makes it highly suitable for integrated platforms. In this work, we demonstrated that 2-3 distinct lifetime species can clearly be discriminated (Fig. 2c and 4d), which opens the door for lifetime-based molecule discrimination as recently used in an integrated protein sequencer^46^.

Furthermore, our scheme is applicable to wide-field super-resolution imaging approaches such as PALM, STORM and DNA-PAINT^47^. The latter would be a natural application of our scheme as binding times can be controlled precisely to yield 1000 photons from ~ 100 ms-long binding events and would yield sub-10% relative error in lifetime considering experimental *F*-values. This opens the door for lifetime-based target multiplexing in super-resolution imaging as demonstrated through confocal FLIM and TCSPC approaches^12,24,48,49^, as well as lifetime-based super-resolved sensing with environment-sensitive probes^50,51^.

## Outlook

This work establishes gated single-photon cameras as a promising tool for spatially multiplexed single-molecule lifetime measurements relying on an optimized scheme enabling time-resolved measurements. The use of this approach should benefit lifetime-based assays of biomolecules for structural biology, diagnostic assays, biopolymer sequencing and single-molecule super-resolution microscopy. SPAD cameras have zero readout noise, which makes them theoretically ideal for high-speed imaging, but two drawbacks remain: warm pixels (Fig. S9) and the lower PDE compared with EMCCD and sCMOS cameras. Nevertheless, we have demonstrated their suitability for single-molecule lifetime imaging, a task that neither EMCCD nor sCMOS cameras could achieve reliably thus far. Compared to confocal TCSPC, our method has a dramatically higher throughput at the cost of a reduced temporal resolution^42^.

Still, we can anticipate several technological improvements in the near future which will improve the detector sensitivity, PDE and decrease the instrument noise, ultimately pushing the bandwidth of our scheme. Improvements in the fill factor and cooling of SPAD cameras are expected to increase the PDE and suppress the warm pixel issue. Newer variants of the SPAD camera that achieve 100% temporal aperture could realize the same scheme without photon rejection^52^, thus decreasing the *F*-values and increasing the temporal resolution by up to 40%. With these combined improvements, we expect our approach to facilitate reliable wide-field single-molecule lifetime measurements at video-rate speeds in the future.

## Materials and methods

### Microscope details

Our setup is a home-built objective-type TIRF microscope based on a commercial inverted microscope body (Olympus IX-71). The light source is an Olympus 12V 100W Halogen Lamp for Olympus U-LH100L-3 Lamphouse.

Illumination was performed with a supercontinuum laser (NKT Photonics SuperK Fianium FIU-15) equipped with a pulse picker to divide the native repetition rate of 78 MHz. A repetition rate of 26 MHz was used. The supercontinuum laser was fiber-coupled to a tunable filter (NKT Photonics SuperK VARIA) which enabled selecting a spectral excitation window of 10 nm or more, centered at the desired wavelength. The unpolarized laser beam coming out of the tunable filter was fiber-coupled to the SuperK collimator, after which the laser beam propagated in free space. A 20× beam expander (Thorlabs GBE20-A) was used in combination with another 2× beam expander (GBE02-A) to flatten the collimated light, which was then focused by an achromatic doublet lens (Thorlabs AC508-300-A-ML) to fill the back focal plane of the high-NA objective (Olympus UPlanSApo100×, NA=1.50). The measured power at the back focal plane was around 5 mW for typical illumination conditions.

The microscope filter wheel is equipped with dichroic mirror/emission filter sets (Semrock Di03-R488-t3-25×36/BLP01-488R-25, Di03-R561-t3-25×36/LP02-561RU-25) that enabled filtering out the excitation light and directing emitted light at the camera. For smFRET experiments, a 600 nm shortpass filter (Thorlabs FESH0600) was placed before the camera to selectively image the donor.

### SPAD camera operation

The camera used in this work is the SPAD512^2^ from Pi Imaging Technology, which is the commercial version of the SwissSPAD2 camera developed by the AQUA group at EPFL^28^. It is a 512 × 512 SPAD array in which the single-photon-sensitive pixels of the camera are operated synchronously. The camera thus outputs binary images, at a maximum speed of 97, 700 frames-per-second (fps). 2^*q*^ − 1 binary images can be binned to reconstruct *q*-bit images akin to conventional EMCCD or sCMOS images, with the difference that no analog-to-digital conversion and no amplification are required, since photons are converted to digital codes directly. Optionally, the camera ‘s time gate can be used to reject photons based on their time delay with a reference signal. It can be precisely positioned by ~ 18 ps steps. It is equipped with a microlens array for a ~ 5-fold increase of the native fill factor of 10.5%^25^. The camera chip was positioned at the primary image plane of the microscope. The built-in optional 1.6× magnification of the microscope was used to achieve a pixel size at the image of 100 nm. An operating voltage *V*_ex_ = 4 V was used for all measurements to minimize noise (at the cost of a slightly reduced PDE^28^). The cooling fans of the camera were used for all measurements. Images were acquired in the 8-bit mode with 20 ms exposure times for proteins in supported lipid bilayer, and 6-bit mode with 10 ms exposure time for DNA origami imaging. Further information on the camera, including a detailed comparison with a standard sCMOS sensor, is available at https://piimaging.com/doc-spad512s.

### Data analysis

#### Acquisition and pre-processing

The camera was operated through the software provided by Pi Imaging technology. 6-bit or 8-bit images were saved in the PNG extension. For processing, images were analyzed using python. The photon counts of pixels were corrected by *(i)* applying the dark count rate mask detailed in Figure S9 to exclude warm pixels and *(ii)* performing pile-up correction.

#### Pile-up correction

Pile-up correction compensates for possible multiple photon arrivals during integration on the single-photon detectors. For this, the measured counts *N*_measured_ were transformed into the actual value *N* used for the analysis in this work using the formula, valid for *q*-bit imaging:

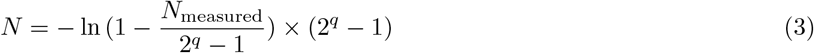

In practice, we only used partial filling of the camera dynamic range (*<*25 photons per pixel per frame for 8-bit images, which is below 10% of the saturation of the pixel), ensuring that the above correction would not distort data much, and would not contribute extra error in lifetime assignments.

#### Molecule detection and intensity trace extraction

Molecules were detected through the Laplacian of Gaussian approach applied to the average image from the first 10 frames following illumination. We consistently used the following parameters: *σ*_min_ = 1, *σ*_max_ = 5, a threshold of 1 and overlap of 0.9 as they worked reliably on our images with a pixel size of 100 nm. The time traces of the molecules were defined as the sum of the corrected photon counts of pixels within a box around the blob center position. We chose the box size to be 5 × 5 pixels (0.25μm^2^) which did not crop emitter spots significantly while minimizing overlap with neighbouring emitters.

#### Lifetime calculations

Following trace extraction, we obtain emitter-wise first gate *N*_*o*_(*t*) and second gate *N*_1_(*t*) time series. We developed two approaches for lifetime extraction: *(i)* a simple, conservative single-lifetime calculation approach for static lifetime measurements, and *(ii)* a more refined, time-resolved lifetime trace calculation. We convolved the trace with a step function and applied a peak finding algorithm to locate intensity change steps. We used a 10-frame duration convolution kernel, and a minimum peak intensity of 15 corrected photons which offered a good compromise between time sensitivity and noise, as shown in Figure S5.

For static lifetime calculations, we only kept stable traces with a single photobleaching step. For the histograms presented in Figure 2c, we applied stringent selection criteria illustrated by the large rejection rate in Figure 2a. The criteria for the time traces are the following: a brightness cutoff to remove the dimmest emitters, a stability criterion to remove high-fluctuation traces, and a single-step photobleaching criterion to ensure that single molecules were imaged. We set the brightness threshold to 50 corrected photons per 20 ms gated frame. Our analysis also identifies the photobleaching time *t*_OFF_. We chose the intensity stability criterion that ON-state fluctuations of *N*_*o*_(*t*) during the interval [0, *t*_OFF_] should not exceed twice the shot noise limit.

For dynamic lifetime calculations, we keep all traces and identify their ascending and descending steps by finding negative and positive peaks in their step-convolved signal. The series of times of ascending and descending steps {*t*_up_} and {*t*_down_} provided a way to sort traces according to their temporal dynamics. We further identified the background level *B* as the lowest observed level following the last photobleaching step. This enabled us to apply equation 1 in a straightforward way to the time trace, which yields a lifetime value per molecule per gate sequence (*N*_*o*_, *N*_1_) (green dots in all plots of Figures S4,S5,S6). As these values were subject to considerable shot-noise-induced fluctuations when sampled at the imaging rate of 50 Hz, we could time-average them to produce more precise results at 25 Hz or 5 Hz (blue dots in the plots). Refining on this approach, we could use the stable segments of the trace identified through convolution and peak finding and provide precise segment-averaged lifetime estimates (black horizontal lines in lifetime plots).

### Sample preparation

#### Aerolysin purification and labeling

The clone of the full-length wild-type aerolysin protein (in the pET22b vector with a C-terminal hexa-histidine-tag (His-tag)) was kindly provided by the Van der Goot laboratory at EPFL. The mutant L249C was generated by Genscript using the wild-type plasmid as a template. Both plasmids were transformed separately into BL21 (DE3 Plys) E. coli cells by heat shock. Cells were grown to an optical density of 0.6 to 0.7 at 600 nm in LB media. Protein expression was induced by the addition of 1 mM IPTG and subsequent growth overnight at 18 °C. Cell pellets were resuspended in lysis buffer (500 mM NaCl, 50 mM tris, pH 7.4), mixed with cOmplete Protease Inhibitor Cocktail, and then lysed by sonication on ice. The turbonuclease solution was added to the resulting suspension and then centrifuged (20,000 g for 30 minutes at 4 °C). After syringe filtration over a 0.45 μm filter, the supernatant was applied to a HisTrap HP column previously equilibrated with lysis buffer. The proteins were eluted as monomers with a gradient of 30 column volumes of elution buffer (500 mM NaCl, 50 mM tris, 500 mM imidazole, pH 7.4). Aerolysin-containing fractions were then buffer-exchanged to the final buffer (20 mM tris, 150 mM NaCl, pH 7.4) using a HiPrep desalting column. The purification of the selected fractions was confirmed by SDS-polyacrylamide gel electrophoresis. The proteins were concentrated to 0.5 mg/mL and stored at −20 °C.

The dyes used for labeling were each dissolved in anhydrous DMSO at a final concentration of 1 mM concentration to ensure stability and stored at −80°C. All steps involving the dyes were performed under light-protected conditions to prevent photodegradation. For the labeling process, mutant L249C was mixed with 10 molar excess of TCEP in a reaction vessel from which air was removed and replaced with an inert gas to prevent oxidation. This mixture was allowed to react for 20 minutes at room temperature. The dye of choice was then added to the reaction mixture at a 10 molar excess compared to the protein concentration. The reaction was left to proceed overnight at 4°C. The protein was purified using 10 kDa cutoff Amicon^®^ Ultra Centrifugal Filter to effectively eliminate the free dye from the sample. A total of five spins (10,000 g for 1 minute each) were performed, with the final buffer added after each spin to ensure the thorough removal of the free dye.

The aerolysin activation process to undergo conformational changes and enable heptamerization includes incubation with trypsin in agarose at a ratio of 1:4 (trypsin:pore, v/v). To ensure that the oligomers consist of one labeled monomer and six unlabeled ones, the labeled monomers of the L249C mutant were mixed with aerolysin wild-type monomers in a molar ratio 1:10, followed by mixing with trypsin and incubating on a spin wheel for 2h at 44 °C. The final labeled proteins were obtained by spinning the sample at 11,000 g for 10 minutes and collecting the supernatant. We ensured elsewhere that the above protocol leads to pore insertion into suspended DPhPC lipid bilayers, which can be monitored by measuring transmembrane ion current^53^. In the next paragraph, we detail the preparation of supported lipid bilayers of the same composition used in this work.

#### Supported lipid bilayer preparation and assembly

DPhPC lipids (0.35-0.4 mg) were initially dissolved in chloroform. The chloroform was removed by subjecting the lipid solution to vacuum evaporation in a desiccator for 1 hour, resulting in a dried lipid film. The film was rehydrated with 1 mL of PBS (pH 7.4) and subjected to brief sonication for 5 minutes to aid in the dissolution of the lipids. The resulting lipid suspension was incubated overnight at 37°C on a heating plate. After incubation, the solution was vortexed until a milky appearance was achieved, followed by further sonication for 1 hour until the solution became clear. To stabilize the vesicles, 1 μL of a 3M calcium chloride stock solution was added to the lipid suspension to reach a final concentration of 3 mM. The vesicle size can be verified using dynamic light scattering. For the layer deposition, coverslips were first cleaned with Hellmanex III detergent to ensure a pristine surface. The deposition chambers, including the coverslip and cylinder, were then plasma-cleaned to enhance the adhesion of the lipid bilayers. The lipid solution was deposited onto the cleaned chamber and incubated in a humidified chamber for at least 90 minutes. Following incubation, the chamber was washed 5-6 times with PBS to remove any unbound lipids and ensure a well-defined lipid bilayer. The proteins were added to the solution at a final concentration of 1 nM and incubated for 10 min, or longer, to reach the desired density of single molecules, before washing again with PBS for another 5 times to reduce the background 10 minutes or longer, to reach the desired density of single molecules, before washing again with PBS for another 5 times to reduce background noise during the measurements. This protocol was adapted from ref.^54^

#### Origami design and assembly

##### DNA origami synthesis

The DNA origami design employed in our studies was designed using CaDNAno6. The structure is available at https://nanobase.org/structure/146 and details can be found in a previous publication^55^. A 7249-nucleotide long scaffold extracted from the M13mp18 bacteriophage (Bayou Biolabs LLC) was folded into the desired shape using 243 staples in 1× TAE (40 mM Tris, 10 mM acetate, 1 mM EDTA), 12 mM MgCl_2_, pH 8 buffer. It was mixed in a 10-fold excess of staples (purchased from IDT) over the scaffold, and 100-fold for the functional staples (fluorophores and biotin, purchased from Biomers GmbH) shown in Table S2. The mixture was heated to 70 °C and cooled to 25 °C at a rate of 1 °C every 20 min. The DNA origami structures were later purified by 1% agarose (LE Agarose, Biozym Scientific GmbH) gel electrophoresis at 70 V for 2 h and stored at 4 °C.

##### DNA origami sample preparation

Coverslips were sonicated in 1% Hellmanex III detergent in deionized water for 10 min, blow-dried with nitrogen and subsequently O_2_ plasma cleaned for 90 seconds. A 100 μm cylindrical glass chamber was then glued to the coverslip using Kwik-kast (World Precision Instruments). A 10 mg/ml BSA-Biotin solution in 1×PBS was then incubated on the surface for 30 min followed by 30 min of 10 mg/ml Neutravidin in 1×PBS to create attachment sites all over the surface of the coverslip. Origamis at a concentration of 50 pM in 1× TAE 12mM MgCl_2_ are added to the surface and their immobilization onto the glass coverslip is monitored during the incubation using a wide-field fluorescence setup. Washing is performed 3 times after each incubation step with 1× TAE 12 mM MgCl_2_.

## Data availability

The data that support the findings of this study will be made available on the Zenodo repository and on EPFL Infoscience repository.

## Author contributions

N.R. and A.R. conceived the project; N.R., A.R., S.B., G.A. and R.R. designed experiments; N.R., S.B. and M.F.M. performed experiments. N.R. analyzed data with input from S.B.; N.R. performed the theoretical analysis and simulations with input from S.B.; M.F.M., M.J.M. and M.D.P. established the protocol for protein construct preparation and control. M.F.M. purified proteins under the supervision of M.J.M., M.D.P. and A.R.; N.S. designed and prepared the DNA origami under the supervision of G.A.; M.F.M., S.B. and N.R. prepared samples; A.R. supervised the project; N.R. wrote the manuscript with input from S.B., R.R., G.A., C.B., E.C. and A.R.; All authors discussed the results and commented on the manuscript.

## Acknowledgements

We acknowledge support from the EPFL Center for Imaging (A.R., N.R., E.C. and C.B.) European Research Council (grant 101020445 to A.R.), the Swiss National Science Foundation (grant 200021-184687 to G.P.A., grant 200021L-212128 to M.D.P. and grant IZSEZ0-224299 to R.R.), the National Center of Competence in Research Bio-Inspired Materials (NCCR 51NF40-182881 to G.P.A.), the European Union Program HORIZON-Pathfinder-Open (grant 101099125 to G.P.A.). We thank Cyril Saudan, Harald Homulle and Ivan Michel Antolovic (Pi Imaging Technology) for their support with the camera software and hardware.

