## Supplementary information for "Wide-field fluorescence lifetime imaging of single molecules with a gated single-photon camera"

| Sample | Dye | Lifetime (ns) | Lifetime standard deviation (ns) |
|---|---|---|---|
| Aerolysin on supported lipid bilayer | LD555 | 1.63 | 0.19 |
| Aerolysin on supported lipid bilayer | Cy3B | 2.29 | 0.19 |
| Aerolysin on supported lipid bilayer | AF488 | 3.34 | 0.18 |
| DNA origami (FRET) | Cy3B | 2.09 | 0.25 |
| DNA origami (no FRET) | Cy3B | 3.45 | 0.24 |

**Table S1**: **Lifetime values reported in this work**, as obtained from gaussian fitting of the lifetime distributions. We find a notable difference between Cy3B in the aerolysin-lipid system and in the DNA origami system. This could be explained by the very different environment of the dye in the two experiments. For example, when bound to single-stranded DNA in DNA-PAINT experiments, an intermediate value of 2.8 ns was found[12].

| Staple Name | Sequence (5'→ 3') |
|---|---|
| Biotin 1 | ACAGGAAGATTGTCCCCCTTATTCACCCTCATTTGTTTC - **Biotin** |
| Biotin 2 | GTTGATAGATATAAGCATAAGTATAGC - **Biotin** |
| Biotin 3 | AGAGTACTCACGCTAACCTTTAATTGC - **Biotin** |
| Biotin 4 | CACTAAAACACTCACGAACTAACACTAAAGT - **Biotin** |
| Biotin 5 | TCACGACGTTGGGCGCTTTGGTAAAAC - **Biotin** |
| Biotin 6 | CAGAGATAGCGATAGTGAATAACATAA - **Biotin** |
| FRET acceptor | ATT ATC ACC GGA AAT GTT AGC AAA CGT AGA A - **Alexa647** |
| FRET donor | AAT ATT GAC GTC ACC GTG CGT AGA TTT TCA GG - **Cy3B** |

**Table S2**: **List of staples for the DNA origami design**. The preparation details are provided in the Methods section.

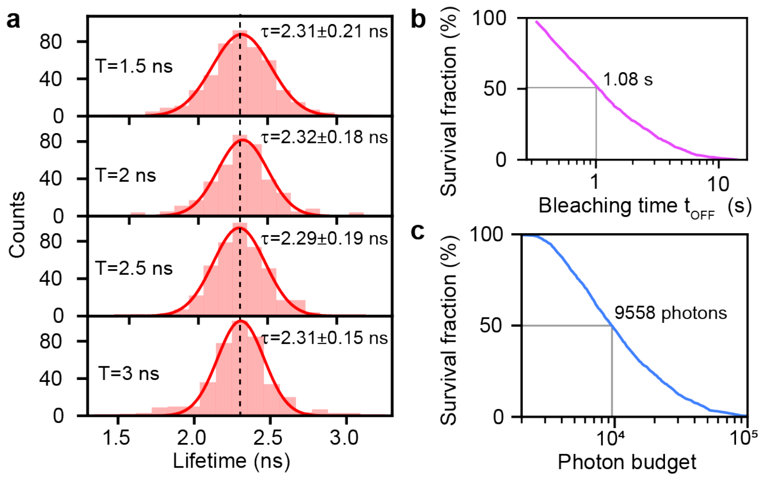


**Fig. S1**: **Extended data on Cy3B-labeled aerolysin in supported lipid bilayer**. **a**, Comparison of single-molecule lifetime distributions with different gate delay $T$. We observe no significant gate delay-induced lifetime bias (dashed line). The distributions contain fewer counts than in Figure 2c as they were taken from a single field of view. **b**, Bleaching time $t_{\mathrm{OFF}}$ survival fraction distribution, showing a median bleaching time of 1.08 s. **c**, Photon budget survival fraction distribution, where the photon budget was obtained by summing all available photons before photobleaching, calculated as $2 \times N_o \times t_{\mathrm{OFF}}/\Delta t$ where $\Delta t$ is the frame integration time. The median photon budget of $\sim 9600$ photons is indicated.

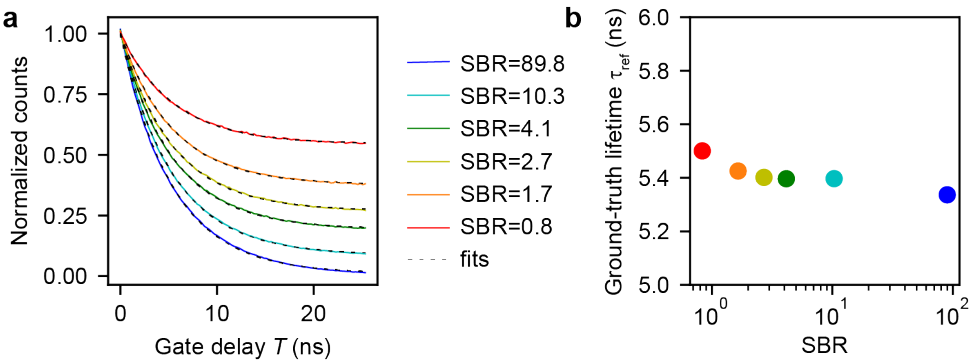


**Fig. S2**: **Determination of reference lifetimes of the fluorescent beads.** Fluorescent beads (PS-Speck™ Microscope Point Source Kit, ThermoFisher Scientific) were imaged at 1% of the power used for single molecules with 3 ms exposure time in 8-bit mode. **a**, The decay was measured using 100 gate delay positions spaced by 256 ps. An average decay was obtained from 90 repetitions, over which no noticeable photobleaching occurred. The reference lifetime $\tau_{\text{ref}}$ was obtained from least-square fitting of the curves to a single exponential with background. **b**, As we found that the background induced mild bias on the lifetime readout, we used the measured reference lifetime value for calculation of the $F$-value curves in Figure 3b. Error bars are smaller than the markers.

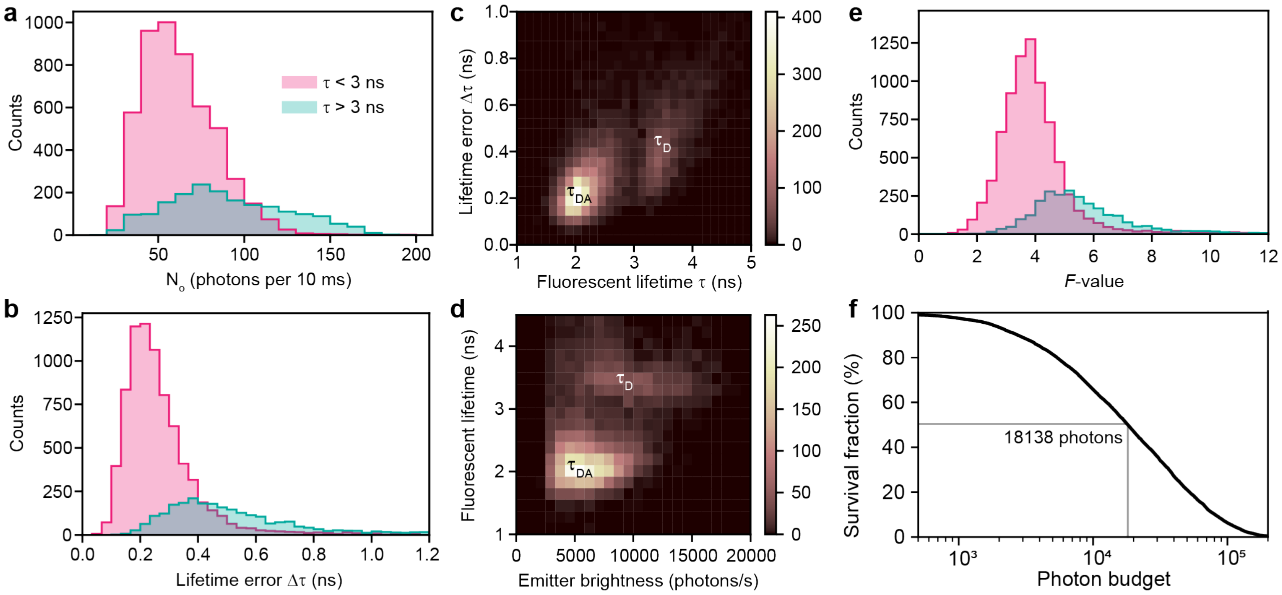


**Fig. S3**: **smFRET extended data underlying Figure 4d. a**, Histogram of $N_o$ values obtained with 10 ms integration time. **b**, Histogram of lifetime errors for 200 ms integration time, calculated as the standard deviation of the 5 Hz sampling trace for traces longer than 600 ms. **c**, Lifetime-lifetime error 2D histogram revealing the two emitter populations. **d**, Brightness-lifetime 2D histogram. The brightness in photons per second is simply $N_o \times 100$. **e**, $F$-value histogram obtained with the lifetime distribution in Figure 4d and the standard deviation in **b**. Equation 2 was applied taking $N = 2 \times 10\, N_o$. **f**, Photon budget cumulative distribution function, limited by photobleaching. The median value of $\sim 18100$ photons is indicated.

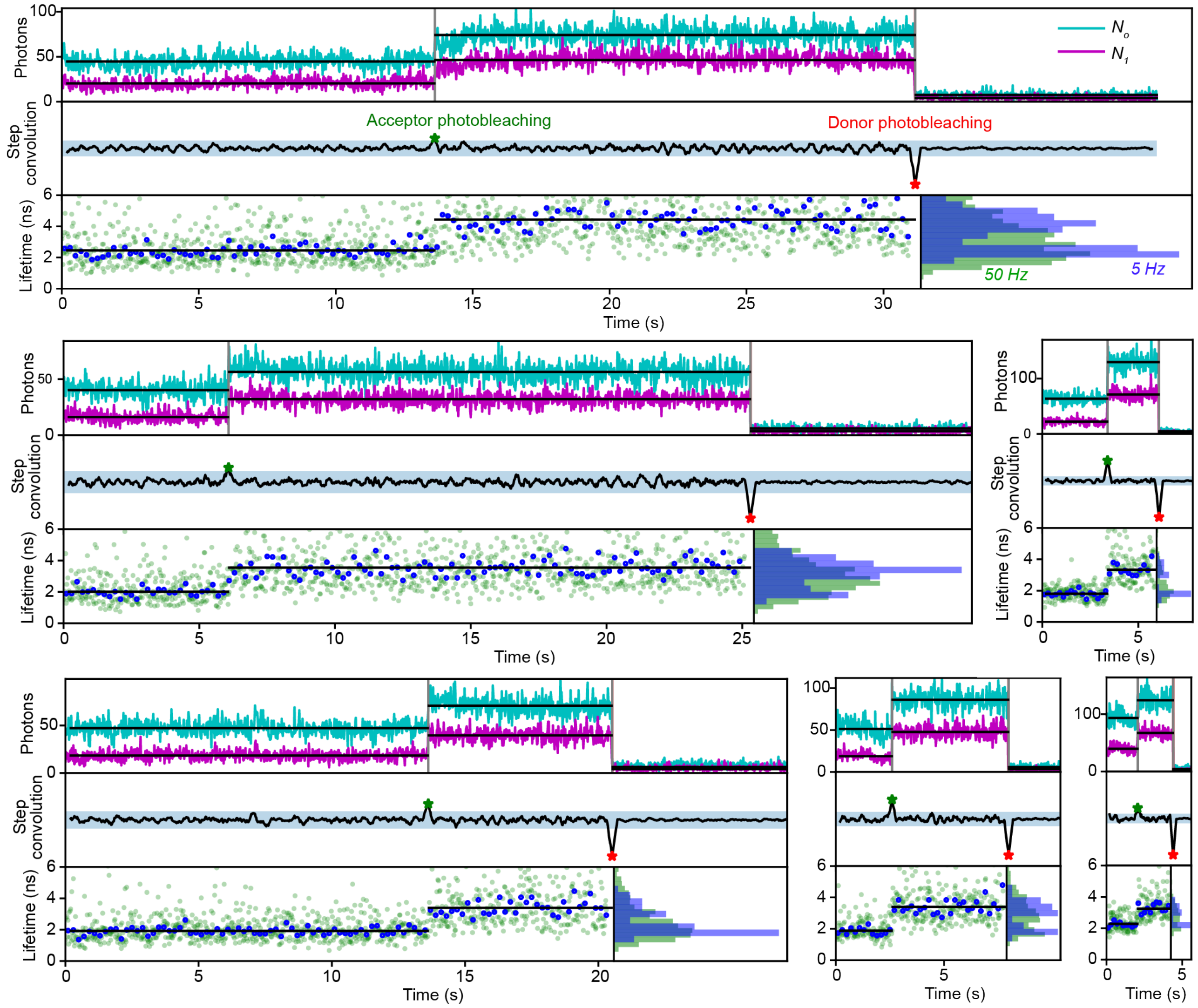


**Fig. S4**: **Donor-acceptor photobleaching representative traces**. Example traces with the raw intensity in the first and second gate (top), its step convolution used for step finding (middle) and the lifetime trace at the native frame rate of 50 Hz and upon ten-fold time binning to obtain a rate of 5 Hz. The trace presented in Figure 4c is reproduced at the bottom left. The highlighted region in the step convolution shows the peak amplitude threshold described in the Methods.

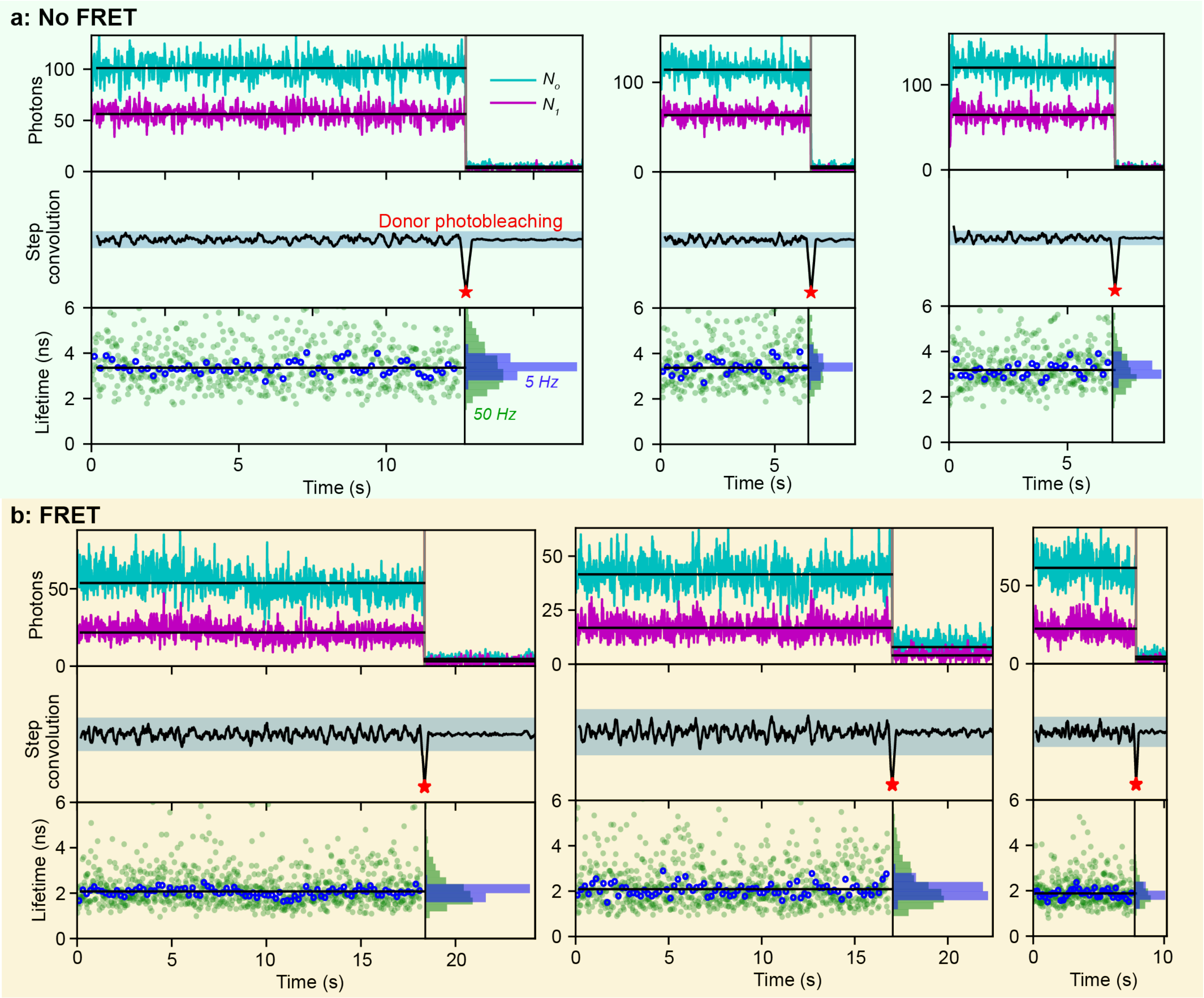


**Fig. S5**: **One-step donor photobleaching traces**. Six example traces of one-step photobleaching traces in the no-FRET (top) and FRET (bottom) cases. Each trace graph layout is the same as in Figure S4. The left traces correspond to the ones provided in Figure 4b.

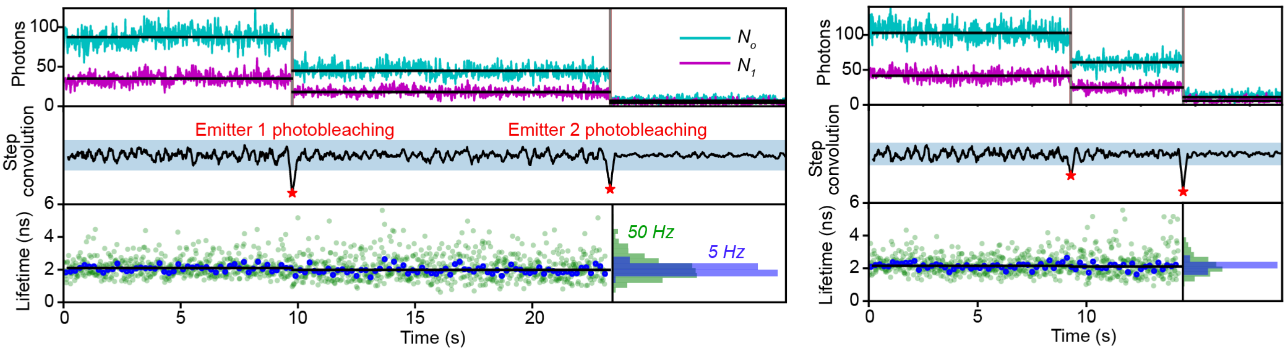


**Fig. S6**: **Two-step donor photobleaching without lifetime change**. Two example traces of two-step donor photobleaching without lifetime change, showing the absence of intensity-induced bias. Each trace graph layout is the same as in Figure S4.

### Derivation of the optimal gate delay

We define the optimal gate delay as the one that minimizes the uncertainty of the lifetime measurement with a given photon budget $N$. The quantity that quantifies the photon cost of the lifetime measurement is the $F$-factor defined by $\Delta\tau/\tau = F/\sqrt{N}$. It is affected by the background level $B$ as well as the choice of the gate delay $T$. It should be noted that some studies use a different definition of the F-value, where $N$ is taken as the number of detected photons rather than the number of available photons, resulting in a metric that does not truly reflect the performance of gated imaging schemes[25]. With the typical values of $N = 100 - 1000$ detected photons used in this work, the pixel-wise binomial statistics of photon counts of the SPAD camera[56] aggregated by emitter point spread function have Gaussian-like statistics and verify $\Delta N \approx \sqrt{N}$.

We extend here the analytical calculation by Heeg[36]: assuming for simplicity a constant non-fluorescent background $B$, the first gate measurement with a fluorescent signal of $S$ reads $N_o = S + B$, and the second gate measurement delayed by $T$ reads $N_1 = Se^{-T/\tau} + B$. Therefore, the lifetime can be retrieved as:

$$\tau/T = 1/\ln\left(\frac{N_o - B}{N_1 - B}\right) = f(N_o, N_1) \tag{4}$$

We thus predict that the error on the $\tau/T$ estimate from $N_o$ and $N_1$ will be given by:

$$\frac{\Delta\tau}{T} = \sqrt{\left(\frac{\partial f}{\partial N_o}\right)^2 \Delta N_o^2 + \left(\frac{\partial f}{\partial N_1}\right)^2 \Delta N_1^2} = \frac{1}{\ln^2(\frac{N_o}{N_1})}\sqrt{\left(\frac{\Delta N_o}{N_o}\right)^2 + \left(\frac{\Delta N_1}{N_1}\right)^2} \tag{5}$$

$$\frac{\Delta\tau}{T} = \frac{1}{\ln^2(\frac{N_o-B}{N_1-B})}\sqrt{\left(\frac{\Delta N_o}{N_o - B}\right)^2 + \left(\frac{\Delta N_1}{N_1 - B}\right)^2} \tag{6}$$

Considering that $N_o \approx S + B$ and $N_1 \approx Se^{-T/\tau} + B$, we obtain:

$$\frac{\Delta\tau}{\tau} = \frac{1}{u}\sqrt{\left(\frac{\sqrt{S+B}}{S}\right)^2 + \left(\frac{\sqrt{Se^{-u}+B}}{Se^{-u}}\right)^2} \tag{7}$$

Where we introduced $u = T/\tau$. Finally, introducing $b = B/S$ (the inverse of the SBR), we obtain:

$$\frac{\Delta\tau}{\tau} = \frac{1}{\sqrt{S}}\sqrt{1 + b + e^u + be^{2u}}/u \tag{8}$$

The estimate of $\tau$ is therefore limited by shot noise (factor $1/\sqrt{S}$), with a prefactor that depends on the choice of the gate delay $T$ with respect to the lifetime of the emitter $\tau$. As the measurement of $(N_o, N_1)$ involved a photon budget of $N = 2(S + B)$, we finally obtain:

$$F(u, b) = \sqrt{2(S+B)}\frac{\Delta\tau}{\tau} = \sqrt{2(1+b)(1+b+e^u+be^{2u})}/u \tag{9}$$

The optimal normalized gate delay $u^*(b)$ is obtained through numerically minimizing $F(u, b)$ with respect to $u$.

### Bias in lifetime estimate

When using equation 1 to estimate the lifetime of an emitter from two gated frames, there can be a bias in the few-photon regime: the distribution of repeated lifetime estimates from N photons $\tau(S, B, T)$ may not have a mean value corresponding to the ground-truth lifetime $\tau_{\text{ref}}$ obtained with $N = S + B \to \infty$ (written as $\tau$ for simplicity in all the above). These can be formalized by considering the average relative bias error $\epsilon(S, B, T) = \frac{\langle\tau(S,B,T)\rangle - \tau_{\text{ref}}}{\tau_{\text{ref}}}$. We can see that this quantify is given by:

$$\epsilon = \frac{\langle f(N_o, N_1)\rangle - f(\langle N_o\rangle, \langle N_1\rangle)}{f(\langle N_o\rangle, \langle N_1\rangle)} \tag{10}$$

A convexity analysis is required to evaluate the bias by first computing second order derivatives.

$$\frac{\partial^2 f}{\partial N_o^2} = \frac{1}{(N_o - B)^2(\log\frac{N_o-B}{N_1-B})^2} + \frac{2}{(N_o - B)^2(\log\frac{N_o-B}{N_1-B})^3} \tag{11}$$

$$\frac{\partial^2 f}{\partial N_1^2} = \frac{-1}{(N_1 - B)^2(\log \frac{N_o - B}{N_1 - B})^2} + \frac{2}{(N_1 - B)^2(\log \frac{N_o - B}{N_1 - B})^3} \tag{12}$$

The bias can be obtained through a second-order expansion of $f$ around $(\langle N_o\rangle, \langle N_1\rangle)$:

$$\epsilon = \Big(\frac{\partial^2 f}{\partial N_o^2}(\langle N_o\rangle, \langle N_1\rangle)\frac{\Delta N_o^2}{2} + \frac{\partial^2 f}{\partial N_1^2}(\langle N_o\rangle, \langle N_1\rangle)\frac{\Delta N_1^2}{2}\Big)/f(\langle N_o\rangle, \langle N_1\rangle) \tag{13}$$

Using the expressions for the derivatives derived above and replacing variables by their mean values, we obtain:

$$\epsilon(S, u, b) = \frac{1}{2Su^2}\big((2+u)(1+b) + (2-u)e^{2u}(e^{-u} + b)\big) \tag{14}$$

As is standard for such statistical estimators, the bias is inversely proportional to the signal.

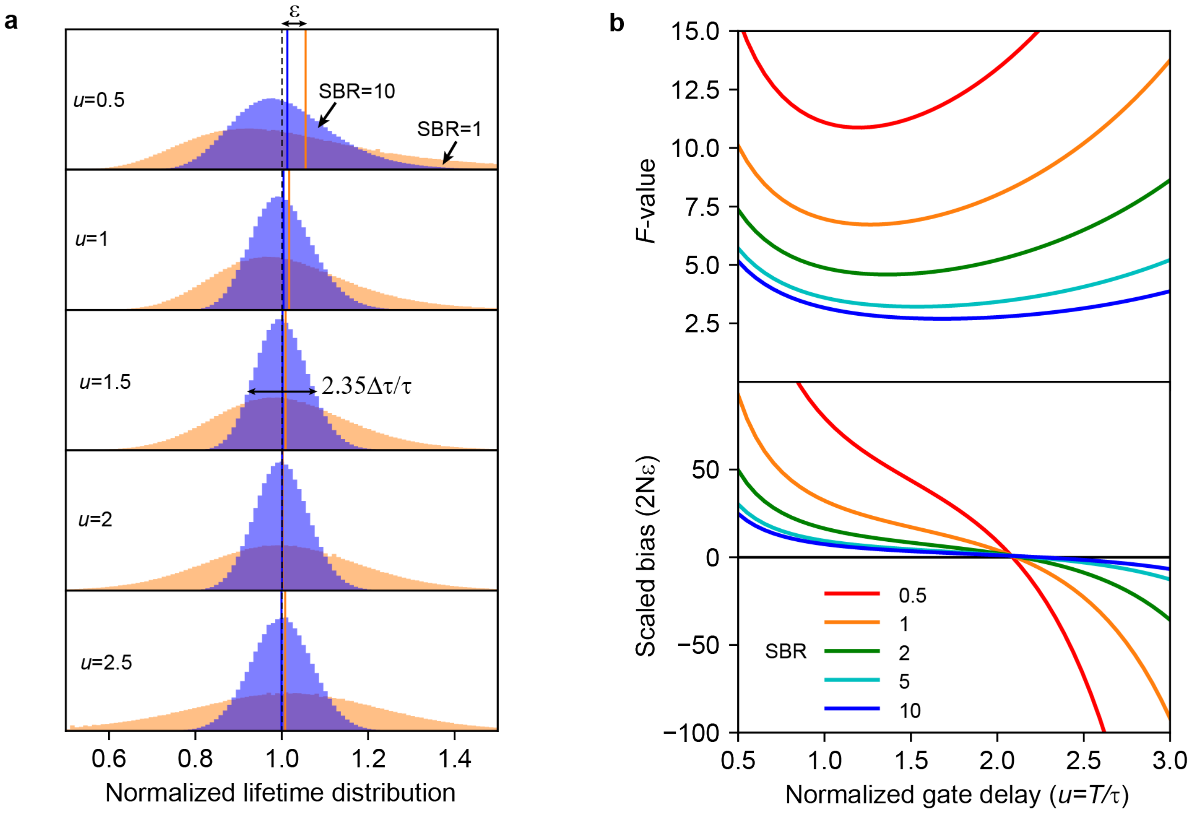


**Fig. S7**: **Variance and bias of the lifetime estimation: Monte Carlo simulations and analytical results.** **a,** Monte Carlo simulations of the lifetime estimate distributions for SBR of 1 (orange) and 10 (blue) with different normalized gate delays $u = 0.5, 1, 1.5, 2, 2.5$. The distribution was obtained by simulating $10^8$ emitters with an average Poissonian photon budget of $2N = 2000$ photons per lifetime estimate. The arrows illustrate both the bias $\epsilon$ and the full width at half maximum of the distribution $\approx 2.35\Delta\tau/\tau$ . **b**, Analytical $F$-value given by $F = \sqrt{2N}\Delta\tau/\tau$ and the scaled bias $2N\epsilon$ as a function of the normalized gate delay for different SBR values ranging from 0.5 to 10. The analytical calculation details above were checked for correctness using Monte Carlo simulations.

### Resolving two lifetimes

A practical application scenario of single-molecule FLIM is to discriminate between two molecules or molecular states based on their lifetimes. We now consider the optimal scheme for resolving two lifetimes $\tau_{\text{short}}$ and $\tau_{\text{long}}$. We define their ratio as $X = \tau_{\text{short}}/\tau_{\text{long}}$ and we aim to optimize the gate delay $T$ to minimize fluctuations of $X$ for the optimal separation of the two lifetimes. For this, we minimize the relative uncertainty on the lifetime ratio given by:

$$\frac{\Delta X}{X} = \sqrt{\frac{\Delta\tau_{\text{short}}^2}{\tau_{\text{short}}^2} + \frac{\Delta\tau_{\text{long}}^2}{\tau_{\text{long}}^2}} \tag{15}$$

### 2-lifetime case

Assuming that the long and short lifetime estimates are given the same photon budget $N$ and have the same background, we obtain:

$$\frac{\Delta X}{X} = \frac{1}{\sqrt{N}}\sqrt{F^2(T/\tau_{\text{short}}, b) + F^2(T/\tau_{\text{long}}, b)} \tag{16}$$

Where $F$ is the function defined in equation 9. Defining $u = T/\tau_{\text{long}}$, this becomes:

$$\frac{\Delta X}{X} = \frac{1}{\sqrt{N}}\sqrt{F^2(u/X, b) + F^2(u, b)} = G(u, b, X) \tag{17}$$

We solve $\frac{\partial G}{\partial u}(u, b, X) = 0$ to obtain the optimal gate position $u^*$ as a function of $X$ and $b$. The results are presented in Figure S8a.

### FRET case

In this scenario, the long lifetime corresponds to the donor lifetime in the absence of FRET $\tau_{\text{donor}}$, and the short lifetime corresponds to the donor lifetime in the presence of FRET $\tau_{\text{FRET}}$ such that the FRET efficiency is given by $E = 1 - \tau_{FRET}/\tau_{\text{donor}}$. The photon budget in this case can be assumed to be $N$ for the donor alone and $(1 - E)N$ for the donor in the presence of FRET. A modified version of equation 17 is then obtained:

$$\frac{\Delta X}{X} = \frac{1}{\sqrt{N}}\sqrt{F^2(\frac{u}{1-E}, \frac{b}{1-E}) + F^2(u, b)} = G_{\text{FRET}}(u, b, E) \tag{18}$$

We solve $\frac{\partial G_{\text{FRET}}}{\partial u}(u, b, X) = 0$ to obtain the optimal gate position $u^*$ as a function of $E$ and $b$. The results are presented in Figure S8b. We find that in both scenarii, for well-separated lifetimes ($X = 0.5$ or equivalently $E = 0.5$) and realistic SBR of $3 - 10$, the optimal value of $u$ is slightly above 1. In other words, to resolve two lifetimes, the optimal strategy is to use a gate delay $T$ close to the longest lifetime $\tau_{\text{long}}$ for the 2-lifetime case or to the donor lifetime $\tau_{\text{donor}}$ in the FRET case. Pathological cases with very high FRET or very low FRET need specific strategies. These calculations can be adapted to cases where the two lifetimes are measured with different SBR and/or photon budgets.

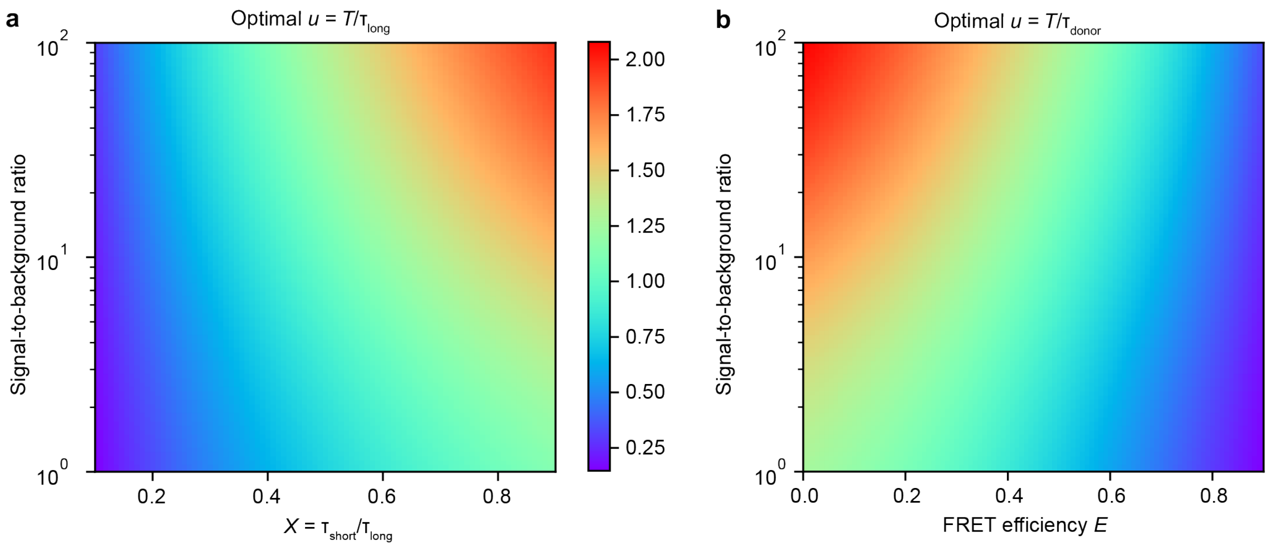


**Fig. S8**: **Resolving two lifetimes.** **a**, 2-lifetime case: optimal gate delay position relative to the long lifetime, as a function of the short-to-long lifetime ratio $X$ and the SBR. **b**, FRET case: optimal gate delay position relative to the donor lifetime, as a function of the FRET effiency $E$ and the SBR. For SBR values in the single-molecule range and $X \approx 0.4$ as in Figure 4, the optimal gate position is close to the long lifetime.

### Camera characterization

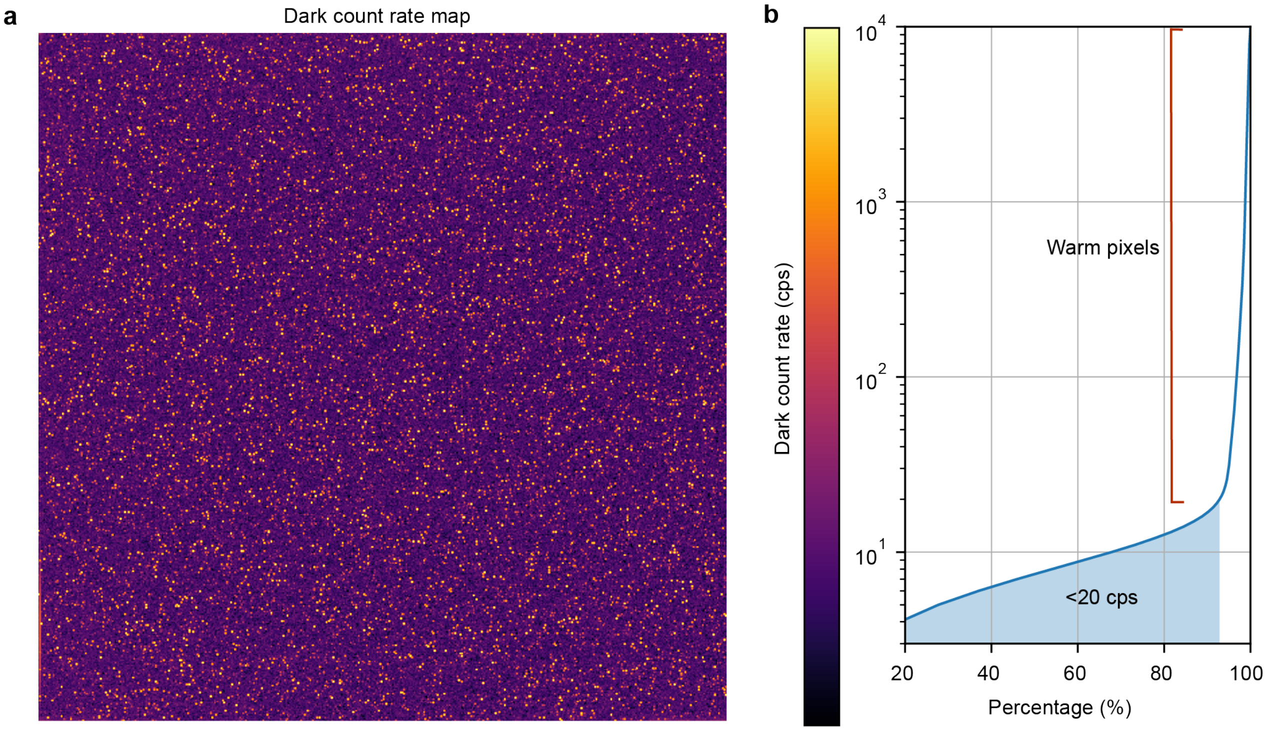


**Fig. S9**: **Dark count rate map and distribution**. **a**, Dark count rate measured by acquiring 10000 frames in the absence of illuminations. Note the logarithmic color scale. **b**, Cumulative distribution corresponding to the map, showing the 5% of warm pixels as the steep ascending part of the curve above 20 counts per second. These pixels were filtered out in the analysis by applying a mask to the images. For more details on the SPAD array noise characteristics, the DCR and the cross-talk of the SwissSPAD3 sensor of similar architecture were recently reported[57].

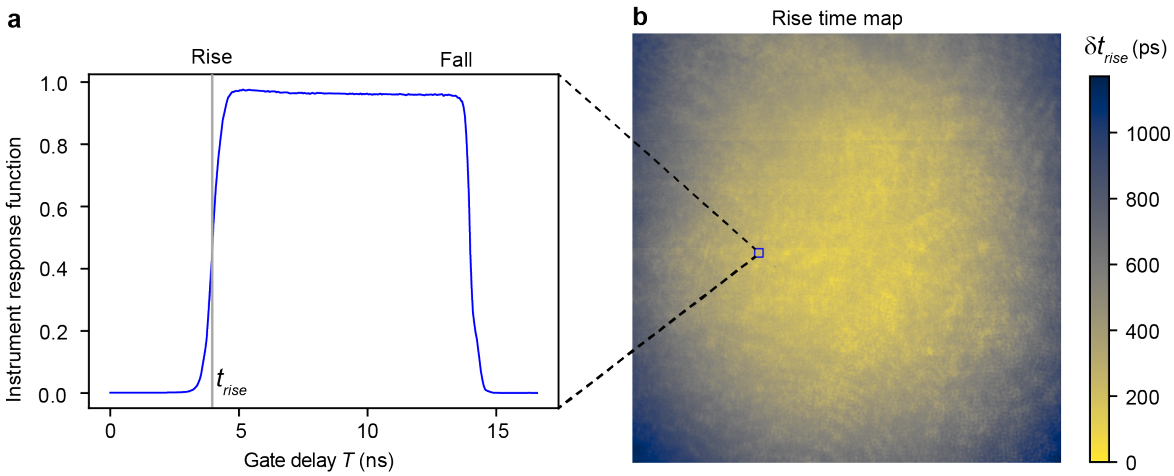


**Fig. S10**: **Instrument response function of the SPAD camera**. **a**, Instrument response function measured as the response to pulsed laser light reflected onto a coverslip, averaged over a $10 \times 10$ pixel region. **b**, Map of the opening time delays showing retardation of the chip edges by close to 1 ns, which was considered in the positioning of the first gate $N_o$. A more complete analysis or the gate properties was done in previous work[28].
